# Differential Analysis of Stromal-Epithelial Interactions between In Situ and Invasive Breast Cancer using Gene Expression Profiling

**DOI:** 10.1101/2022.07.30.502150

**Authors:** Adam Officer, Andre M. Dempsey, Lyndsay M. Murrow, Zev Gartner, Pablo Tamayo, Christina Yau, Olivier Harismendy

## Abstract

**Background:** Changes in microenvironment cell-cell interactions (CCI) during the progression from ductal carcinoma in situ (DCIS) to invasive ductal carcinoma (IDC) are poorly understood. Gene expression studies are confounded by cellular heterogeneity and few separate stromal and epithelial contributions, resulting in a lack of reliable prognostic biomarker to guide treatment decisions.

**Methods:** The gene expression of 293 microdissected regions from DCIS (92 epithelial, 31 stromal) and IDC (78 epithelial, 30 stromal) cases was aggregated from 6 datasets. Expression signatures of 6 cell lineages extracted from normal breast single-cell profiling were used to correct for differences in cell abundance. Subtype-specific functional differences between DCIS and IDC were measured for each region type using Gene Set Enrichment Analysis (GSEA). DCIS-IDC stromal-epithelial interactions were compared using the expression product of 139 ligand-receptor (LR) pairs permuting the DCIS-IDC labels to assess significance.

**Results:** Variation in cell-lineage abundance separated epithelial regions into 4 clusters, including one enriched for DCIS (Myoepi-Enriched) and two for IDC (Infiltrated, Vascularized). GSEA on cell lineage normalized expression data identified subtype-independent changes in epithelial regions (induction of Extracellular Matrix maintenance genes, reduction of Tp53 signaling in IDC), as well as subtype-specific changes (proliferation in ER- and Her2-IDC, reduction in Nucleotide Excision Repair in ER+ IDC). In the stroma, Notch and Rho-GTPase signaling were induced in IDC irrespective of subtype. The stromal-epithelial interaction level of 6 and 4 LR pairs were significantly enriched in DCIS and IDC, respectively. Five of the 6 DCIS-enriched LR pairs involved ephrin interactions, with interaction level progressively decreasing from normal to DCIS to IDC. In contrast, 2 IDC-enriched LR pairs involved T-cell activity likely regulating Treg proliferation (*CD28-CD86*) or T and NK cells stimulation (*CD226-PVR*). Notably, the bulk expression product of one identified LR pair (*EPHB4-EFNB1*) was associated with poor survival in IDC (HR=1.47, p=0.04) suggesting that early remodeling of this stromal-epithelial interaction may have long-lasting impact on disease severity.

**Conclusions:** The observed changes in cell states and stromal-epithelial interactions, beyond those driven by difference in cell abundance, may lead to new biomarkers for prognosis and targets for secondary prevention.

## Introduction

Ductal carcinoma in situ (DCIS) is considered a non-obligate precursor to invasive ductal carcinoma (IDC), but the mechanisms driving invasion are not fully understood. Observational studies suggest that a large fraction of DCIS lesions may never progress to IDC^1,2^. Multiple genomic and transcriptomic studies have investigated the molecular differences between DCIS and IDC and highlighted key differences and similarities between these two diseases. Prior work has shown that DCIS synchronous with IDC lesions harbor similar copy number alterations (CNAs) and mutations.^3^ Gene expression profiling of DCIS and IDC has identified similar molecular subtypes in both stages suggesting similarities between the premalignant and malignant stage^4^. Additionally, there are small gene expression changes between DCIS and IDC with only a handful of significantly differential genes identified across several studies.^5,6^

Among the gene expression differences identified, many involve signaling pathways participating in intercellular communication.^7,8^ Within the epithelial compartment, extracellular matrix and cell adhesion processes are significantly changing between DCIS and IDC, suggesting that the surrounding intercellular space is being restructured. Several studies of microdissected histological regions suggest that more gene expression changes take place in the surrounding stroma as compared to malignant epithelial cells^5^. Follow-up studies have repeatedly identified changes in signaling pathways involved in oncogenesis and invasion, including extracellular matrix, cell adhesion and intercellular communication.^5,9^ These results have somewhat been inconsistent, although not necessarily conflicting. Such gene expression micro-environmental studies have remained small and hard to validate due to the small size, systemic archival of DCIS tissue, and the labor required for microdissection, hampering the generalization and further exploration of the findings.

With recent advances in methodologies and analytical tools, it is now possible to measure microenvironment signaling, specifically through cell-cell interactions (CCI), using gene expression from single-cell RNA-seq as well as bulk gene expression.^10,11^ In general, CCI methods quantify coordinated expression between known ligand and receptor pairs and compare these values to a background model in order to identify significantly active CCI. However, the vast majority of these methods are not able to compare CCI between two different groups, in this case DCIS and IDC, to identify differential CCIs that may be mediating phenotypic changes accompanying breast cancer progression.

Here we present the results of a meta-analysis that compares stromal-epithelial CCI between DCIS and IDC using publicly available gene expression studies of micro-dissected stromal and epithelial regions from 163 patients. We characterize differences in gene expression and cell-cell interactions occurring at the interface between stromal and epithelial compartment, highlighting the contribution of microenvironment factors to breast cancer invasion, validating previous findings, prioritizing ligand-receptor pairs in their micro-environment context and assessing their contribution to overall survival.

## Results

A large gene expression dataset was assembled from 6 independent gene expression studies and merged after correcting for batch effects (see **Methods**). A total of 293 distinct gene expression profiles from epithelial and stroma regions were identified, including 46 from normal biopsies, 123 from DCIS, 108 from IDC and 16 from other breast malignancies representing a total of 163 distinct patients (**Table S1**). Using principal component analysis, samples corresponding to stromal and epithelial regions are well separated while study of origin cannot be distinguished (**Figure 1A,B**), therefore indicating the aggregation of the data was effective and did not introduce spurious variation. Furthermore, hierarchical clustering suggests that within both stromal and epithelial compartments the normal regions are distinct from non-normal regions with small differences between DCIS and IDC observed (**Figure 1C,D**).

**Figure 1:**
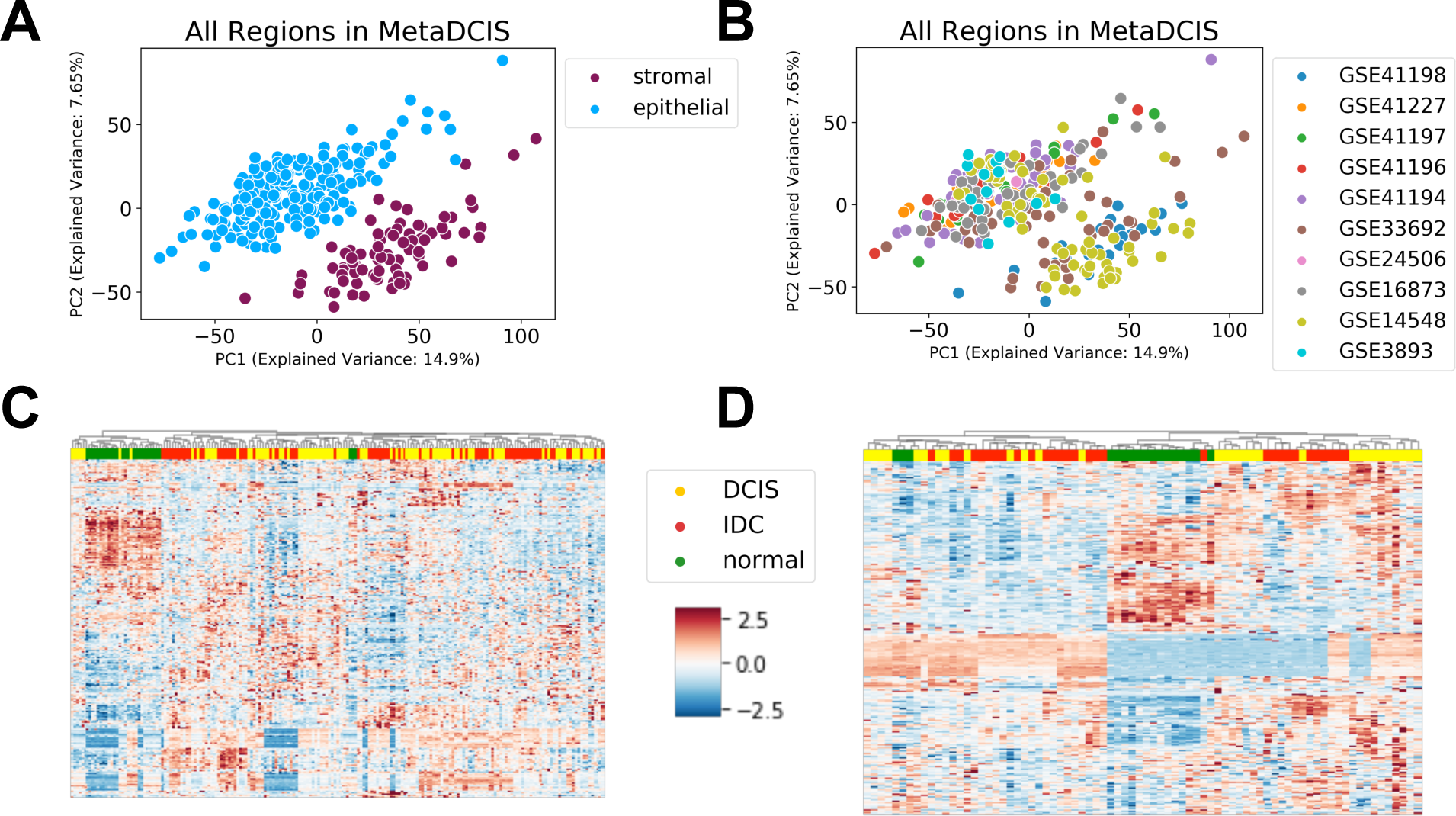
Unsupervised classification of gene expression profiles. **(A-B)** Principal Component Analysis (PCA) of all gene expression datasets included in the study after batch-effect correction and colored by tisssu compartment (A) or source dataset (B). **(C-D)** Hierarchical clustering of the epithelial © or stromal (D) expression profiles using the expression of the 500 most variable genes.

Gene expression signatures were used to classify epithelial regions by intrinsic subtypes (PAM50 - **Table S1**). Her2 and ER status was determined by expression of ERBB2 and ESR1, respectively (see **Methods**). We observed the expected enrichment of ER samples in Luminal subtypes, Her2 enrichment in Her2-like subtype and high prevalence of ER-/Her2-in Basal-like subtypes, indicating the validity of the expression based subtyping. Luminal A was more frequently observed in DCIS lesions (21/30, p=1.7*10^−3^, Fisher exact test) and Luminal B was more frequently observed in IDC lesions (38/59, p=5.6*10^−5^, Fisher exact test). Furthermore, there was a trend towards the enrichment of the Her2-like subtype in DCIS samples (17/92, p=0.045, Fisher exact test). Hence, while the original studies were not designed to faithfully represent a breast oncology clinic, the composition of the aggregated cohort was broadly consistent with previous studies^4^ and likely captured the histological and molecular diversity of breast DCIS and IDC.

### Changes in Stromal and Epithelial Cellular Composition

To explore the role of the tumor microenvironment in breast cancer invasion, we first used the bulk expression profiles to estimate and compare the abundance of multiple cell lineages in both the epithelial and stromal compartments with CIBERSORTx (see **Methods**). Computational deconvolution of cell types by CIBERSORTx works optimally when provided with high-quality cell lineage reference gene expression profiles, ideally derived from large clusters of single-cell datasets. With no large-scale DCIS single-cell gene expression data publicly available and the risk of skewing the results for IDC biology, we chose instead to use the profiles of 86,136 cells observed in normal breast tissue specimens obtained from 28 breast reduction mammoplasty donors. This single-cell dataset consists of a breadth of both stromal and epithelial cell types, is of high data quality and has been fully characterized^12^. The cell type clusters were aggregated into 6 broad cell lineages - luminal, myoepithelial, macrophages, lymphocytes,endothelial and fibroblasts - which were used to estimate cell abundance in normal, DCIS, and IDC stromal and epithelial regions (**Table S2**).

Hierarchical clustering of the epithelial samples based on cell lineage abundance estimates identified four different clusters labeled according to their dominant lineage: Myoepi-Enriched, Infiltrated, Vascularized and Luminal-Enriched (**Figure 2A**). The Infiltrated and Vascularized clusters were enriched for IDC samples (16/78, p=0.01 and 17/78 p=0.018, respectively, Fisher Exact Test, **Figure 2B**). In contrast, the Myoepi-Enriched cluster was enriched in DCIS samples (28/92 p=0.009, **Figure 2B**). These observations suggest that epithelial IDC samples have proportionally more lymphocytes, macrophages and endothelial cells and fewer myoepithelial cells. A supervised univariate approach corroborated these results (**Figure 2C**) and further showed that fibroblasts are 67% more abundant in IDC compared to DCIS (FDR<0.1).

**Figure 2:**
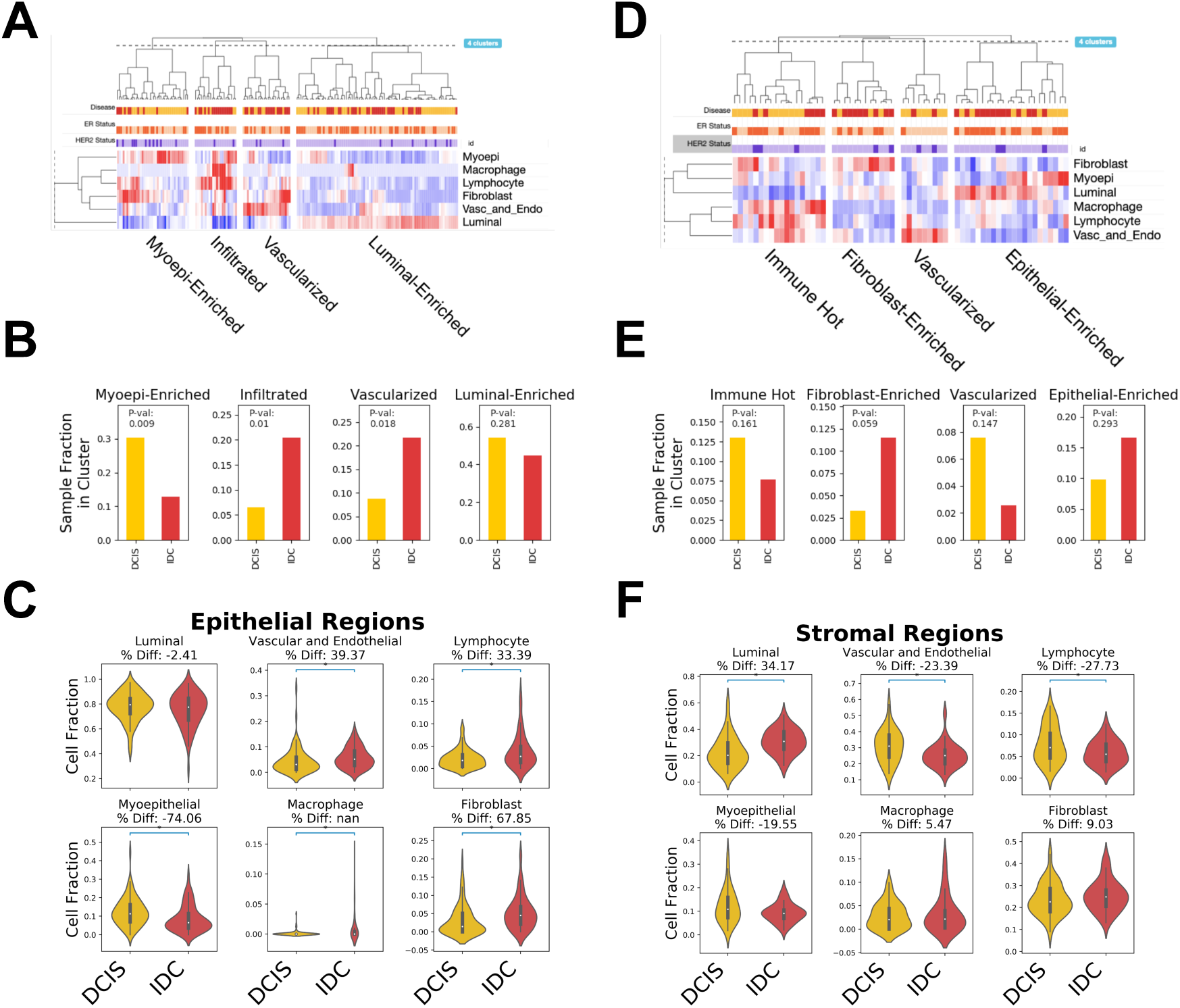
DCIS-IDC expression differences associated with cellular composition. **(A,D)**. Hierarchical clustering of epithelial (A) or stromal (D) regions according to the prevalence of 6 cell-lineages (color-scale - z-scores) as measured using CIBERSORTx. Clusters are labelled according to the dominant pattern in cellular composition. **(B,E)** Prevalence of DCIS (yellow) and IDC (red) region in each cluster for epithelial (B) or stromal (E) regions. P-values were obtained using Fisher Exact test. **(C,F)** Univariate comparison of fraction of each cell lineage between DCIS and IDC in epithelial (C) and stromal (F) regions.

Similarly, hierarchical clustering of the stromal samples based on cell lineage abundance estimates subset of the cell abundance matrix identified four different clusters: Immune Hot, Fibroblast-Enriched, Vascularized and Epithelial-Enriched (**Figure 2D**). None of the clusters showed a significant enrichment between DCIS and IDC stromal regions, to the exception of a trend toward higher fraction of IDC in Fibroblast-Enriched stroma (9/30 p=0.059 Fisher Exact test, **Figure 2E**). Univariate analysis identified a median of 34% more luminal cells and 27% and 23% fewer lymphocytes and vascular and endothelial cells in the stroma adjacent to IDC regions (**Figure 2F**). This may be due to contamination from imperfect microdissection, as previously reported^5^, or imperfect cell lineage annotation by the CIBERSORTx algorithm^13^. Importantly, the important differences in cellular composition of DCIS and IDC identified can be a major confounding factor when trying to use bulk expression profiling to investigate molecular changes associated with invasion.

### Stromal and epithelial functional differences between in situ and invasive lesions

In order to explore DCIS-IDC molecular differences beyond differences in cellular composition, we corrected the bulk gene expression matrix by relying on the linear properties of the CIBERSORTx method. Specifically, we derived a cell abundance corrected gene expression matrix from a normalized cell abundance matrix and the cell specific expression tensor (see **Methods**). The resulting corrected gene expression matrix was then used to investigate functional differences using gene set enrichment analysis (GSEA).

We first compared epithelial regions. The large size of the assembled cohort allowed us to split the analysis according to ER and HER2 status. We compared the enrichment for 803 gene sets (Cancer Hallmark and Reactome from MSigDB^14^) across four subtypes of epithelial regions: ER+ (40 IDC samples, 41 DCIS samples), ER- (51 IDC, 38 DCIS), Her2+ (7 IDC, 17 DCIS), Her2- (71 IDC, 75 DCIS), recognizing the overlaps between ER and Her2 classifications. There were 215 gene sets whose expression was significantly altered (FDR<0.01) in at least one of the comparisons and these gene sets grouped according to 4 enrichment patterns (**Figure 3A, Table S3**) : 1) *Enriched in IDC* included gene sets related to extracellular matrix organization and degradation which likely reflect how the epithelial cells actively participate in the degradation of the matrix in invasive disease; 2) *Depleted in IDC* included gene sets such as regulation of Tp53 activity, expression of ion and vitamin transporters or G protein signaling likely reflecting the stronger reliance of invasive disease on the loss of G1 checkpoint and changes in cellular metabolism and environment sensing; 3) *Enriched in IDC in ER- and Her2- context* - the largest group - included gene sets related to proliferation and cell cycle control, indicating strong proliferative differences from DCIS to IDC in triple negative cancer which represent the majority of this category; and finally 4) *Depleted in ER+ IDC*, were genesets associated with Nucleotide Excision Repair, targets of Myc transcription factors and mitochondrial protein import suggesting that in the context of ER signaling, these processes were dispensable, or perhaps detrimental, to IDC growth.

**Figure 3:**
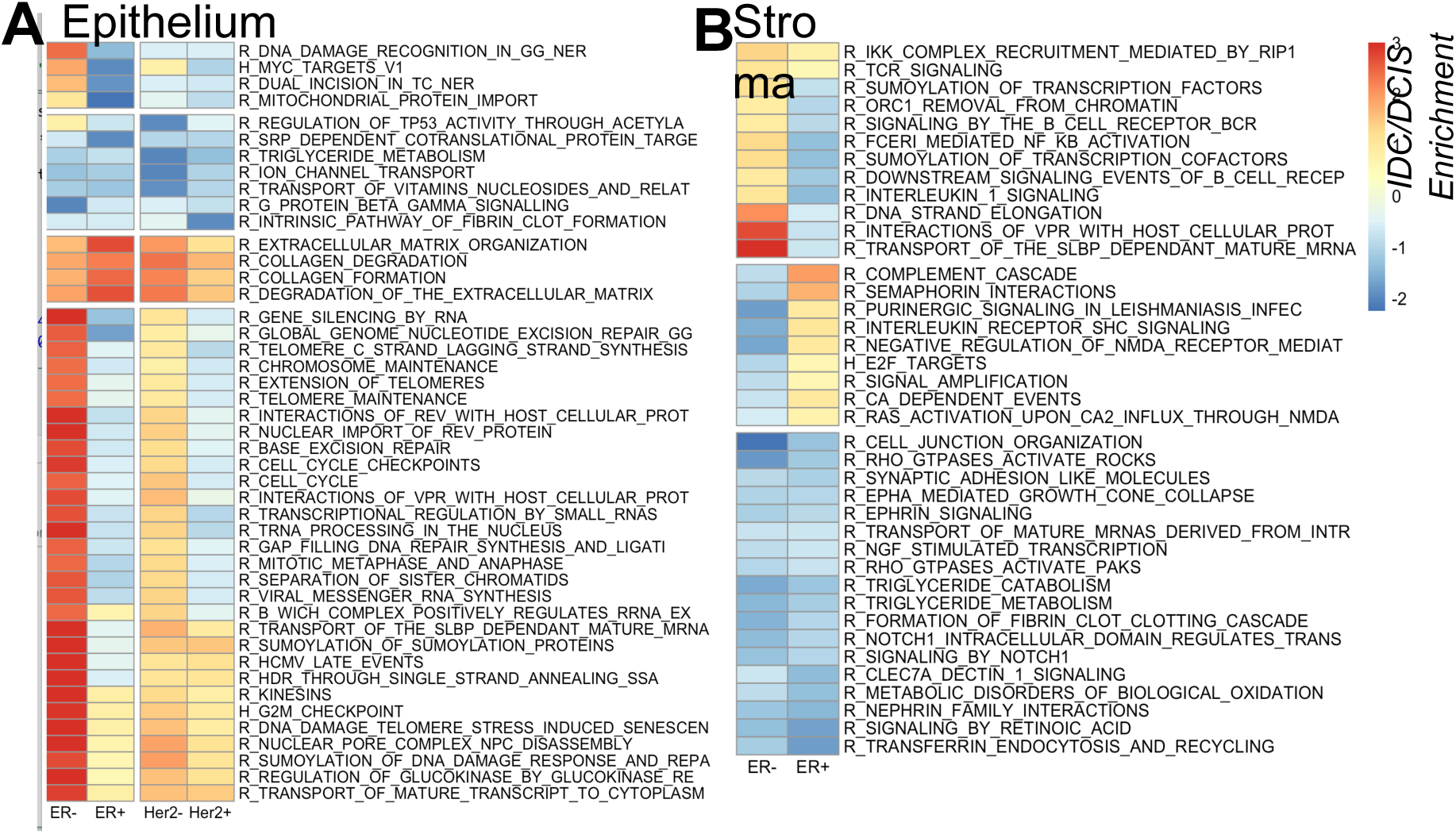
Context specific functional differences between DCIS and IDC. **(A-B)** Heatmap of GSEA normalized enrichment scores (positive/red: enriched in IDC, negative/blue: enriched in DCIS) from the differential gene expression between DCIS and IDC epithelial (A) or stromal (B) samples from different subtypes (columns, ER and Her2 overlap). Gene sets significant (FDR<0.05) in at least one context and with highest absolute enrichment scores are displayed (epithelium: 2.4, stromal: 1.8). Gene set names are truncated and/or abbreviated (R: Reactome, H: Hallmarks) for visualization clarity. Full results in Table S3.

In the stromal region a similar analysis was conducted albeit limited to 2 subtypes: ER+ (17 IDC, 14 DCIS) and ER- (13 IDC, 17 DCIS) with sufficient number of samples (**Figure 3B**). There were 167 gene sets whose expression was significantly altered in one or both comparisons (FDR<0.01). Notably, after correcting for cell type abundance differences, the differences observed reflect phenotypic changes in the stroma, or cell abundance differences not accounted for (e.g. B vs T lymphocytes). The significant gene sets included increased TCR, BCR and Interleukin signaling in the ER- context, likely reflecting an immunoactive state, in addition to the known higher infiltration. In the ER+ context, semaphorin and complement signaling were enriched in IDC. Finally, a number of pathways were depleted in IDC irrespective of the ER status. These processes are involved in the remodeling of cell-cell communication and adhesion - notably with Cancer Associated Fibroblasts - and mediated by signaling through Ephrin, Rho GTPase, C-type lectin or Notch1, all previously associated with breast cancer invasion. These pathways may therefore be dysregulated in fibroblasts in reaction to the invasion, and/or in the residual epithelial cells remaining at the invasive front.

The results generally validate previous gene expression studies. The size of the assembled cohort, combined with separation of epithelial and stromal regions allows a more refined perspective, especially distinguishing the relative importance of processes in different histological subtypes. In turn, the correction for cell-type abundance enabled the specific assessment of phenotypic signaling differences in cells in the micro-environment, pinpointing to the importance of interactions with fibroblasts in invasion.

### Differences in Stromal-Epithelial Interactions between DCIS and IDC

Gene set enrichment analysis highlighted important differences in the functional states of the DCIS and IDC epithelial and stromal regions, but did not capture possible synergies or antagonism between them and how interaction between the malignant and non-malignant compartments can drive or repress invasion. We hypothesize that several of the changes observed are triggered by ligand-receptor interactions occurring between cellular compartments or between a cell and its environment. To investigate this, we selected 139 pairs of ligands and receptors from CellPhoneDB (**Methods**)^15^ and used their gene expression level to estimate the strength of their stromal-epithelial interaction in both IDC and DCIS contexts. Each known ligand-receptor pair was tested in both directions: epithelial ligand and stromal receptor (ES) and the reciprocal epithelial receptor and stromal ligand (SE), for a total of 278 interactions measured. Each interaction was estimated separately in DCIS and IDC context and significant differences were assessed using an empirical permutation strategy of the DCIS and IDC labels to compute the false discovery rate (**Methods**).

Using this framework a total of 10 interactions, involving 8 ligands and 8 receptors significantly different (FDR<0.1) between DCIS and IDC (**Figure 4A, Table S4**). Six interactions were stronger in DCIS and all involved ephrins, themselves both ligands and receptors depending on the context^16^. Ephrins participate in cell localization, guidance and compartmentalization of tissues and have previously been implicated in breast cancer progression and invasiveness^17^. Of the 4 interactions estimated to be stronger in IDC, the *WNT2-FZD2* interaction was the strongest in both the ES and SE directions (normalized difference in interaction score of -0.88 and -0.63, respectively, **Figure 4A**). This SE finding is consistent with the role of Fzd2 and Wnt signaling in mediating EMT, invasion and metastasis in cancer 18^,19^. Furthermore, 2 out of 4 interactions stronger in IDC involved immune-receptors: *CD86-CD28* (ES), is a co-stimulatory interaction required for TCR activation by T cells^20^, while *CD226-PVR* (ES), is an interaction known to sensitize NK and T-cell mediated lysis of tumor cells^21^. Given that the expression values were corrected to account for variation in cellular composition, these interaction changes may reflect a change of state of the cell in a particular histological region, or can be the consequences of sub-lineage differences not accounted for (T vs B lymphocytes).

**Figure 4:**
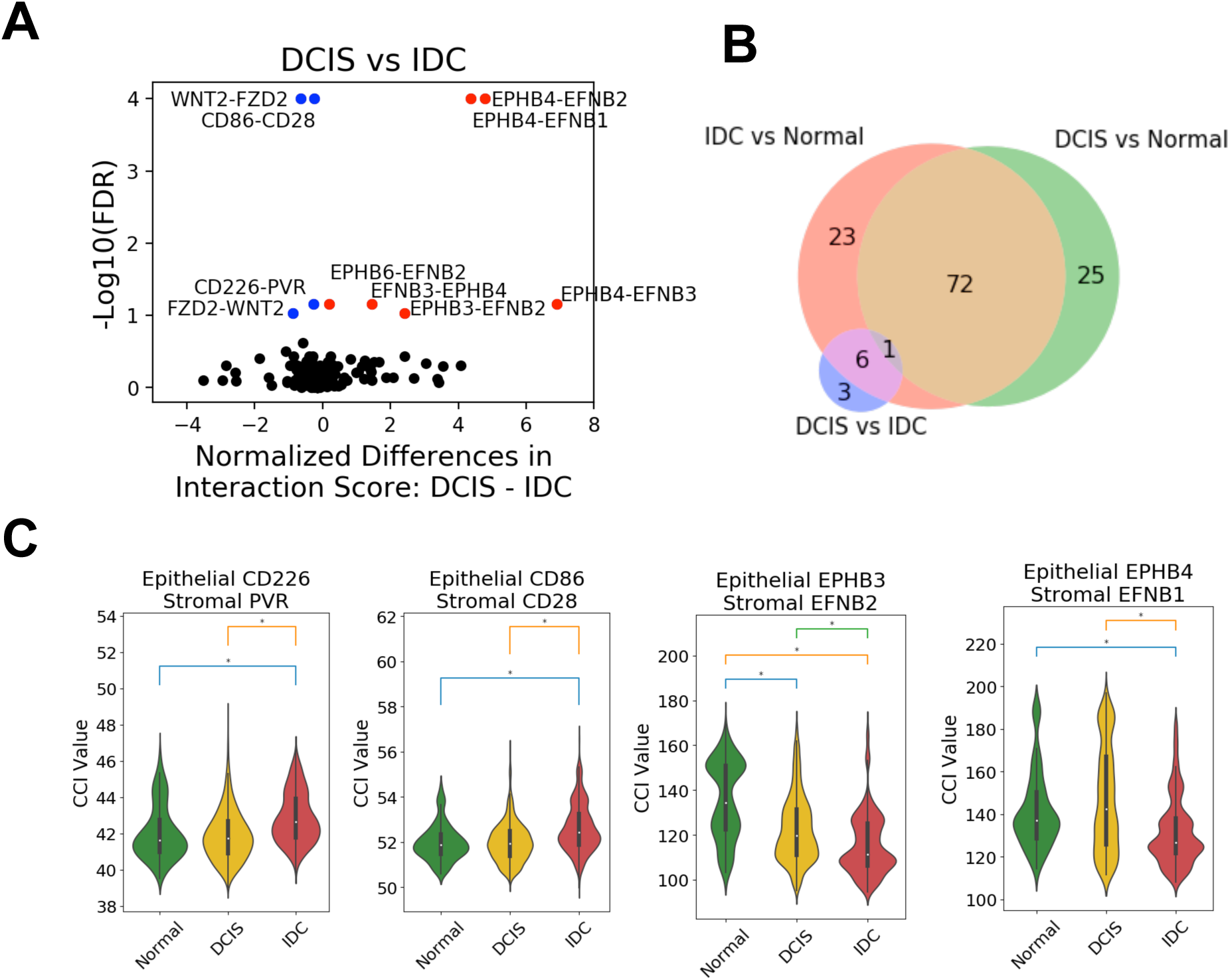
Differences in stromal-epithelial cell-cell interactions. **(A)** Differences in DCIS-IDC interaction scores (x-axis) for all tested LR pairs (points) in both SE and ES directions. Significance was estimated using False Discovery Rate (y-axis) and significant LR pairs stronger in DCIS (red) or IDC (blue) are labelled. Full results in Table S4). **(B)** Venn-diagram displaying the number and overlap of significantly different interactions observed in DCIS-IDC, DCIS-normal and IDC-normal comparisons. **(C)** Distribution of the stromal-epithelial LR Interaction scores (y-axis) measured in normal (green), DCIS (orange) and IDC (red) context for 4 selected LR pairs significant in the DCIS-IDC comparison. (*) indicate FDR<0.1.

In order to understand at which point in breast carcinogenesis these interactions shift we used the adjacent normal stromal and epithelial regions as an outgroup. By comparing the strength of the interactions between normal and DCIS or normal and IDC we can determine how early in oncogenesis such interactions are being remodeled. Of the 278 interactions tested, there were 98 and 102 interactions that are identified as significantly changing between normal-DCIS and normal-IDC, respectively. This is an order of magnitude more interactions than those changing in the DCIS-IDC comparison (**Figure 4B**). This suggests that the CCI remodeling occurs early in oncogenesis, prior to invasion. This was the case for all of the ephrin-mediated interactions that showed a monotonic decrease from normal to DCIS to IDC suggesting that these interactions are progressively changing between both the normal to DCIS transition as well as the DCIS to IDC transition (**Figure 4C**). In contrast, the *CD226-PVR* (ES) interaction was identified as significantly stronger in IDC compared to either DCIS or normal, timing this interaction to change with the DCIS to IDC transition. A similar observation was made for *CD86-CD28* (ES), indicating that the activation of immune-related CCI is a later event in the disease progression.

### Association with Breast Cancer Survival

The changes of CCI between DCIS and IDC, while identified from the microdissection of local disease, may also mediate metastatic spread and have important consequences on survival. We analyzed The Cancer Genome Atlas (TCGA) to determine, within an IDC cohort, if these interactions had prognostic significance. In absence of microdissection, we approximated ES and SE CCI scores using the bulk expression profile (**Methods**). To account for known prognostic factors such as ER positivity, tumor stage, advanced age, immune infiltration, we used Cox proportional hazards (CoxPH) regression to determine the prognostic value of candidate CCI score (**Figure 5A**). Of the 10 ligand-receptor interactions that were significantly different between DCIS and IDC, we identified one interaction, EPHB4-EFNB1, that was significantly associated with survival (HR=1.47, p=0.04, **Figure 5B**) and the 3 year survival of patients patient with high *EPHB4-EFNB1* expression product was worse (76% vs 88% - **Figure 5C**). Notably the interaction between the two is required as neither the expression of *EPHB4* nor *EFNB1* alone significantly affected survival (p=0.18 and 0.34, respectively, **Figure 5D,E,F**).

**Figure 5:**
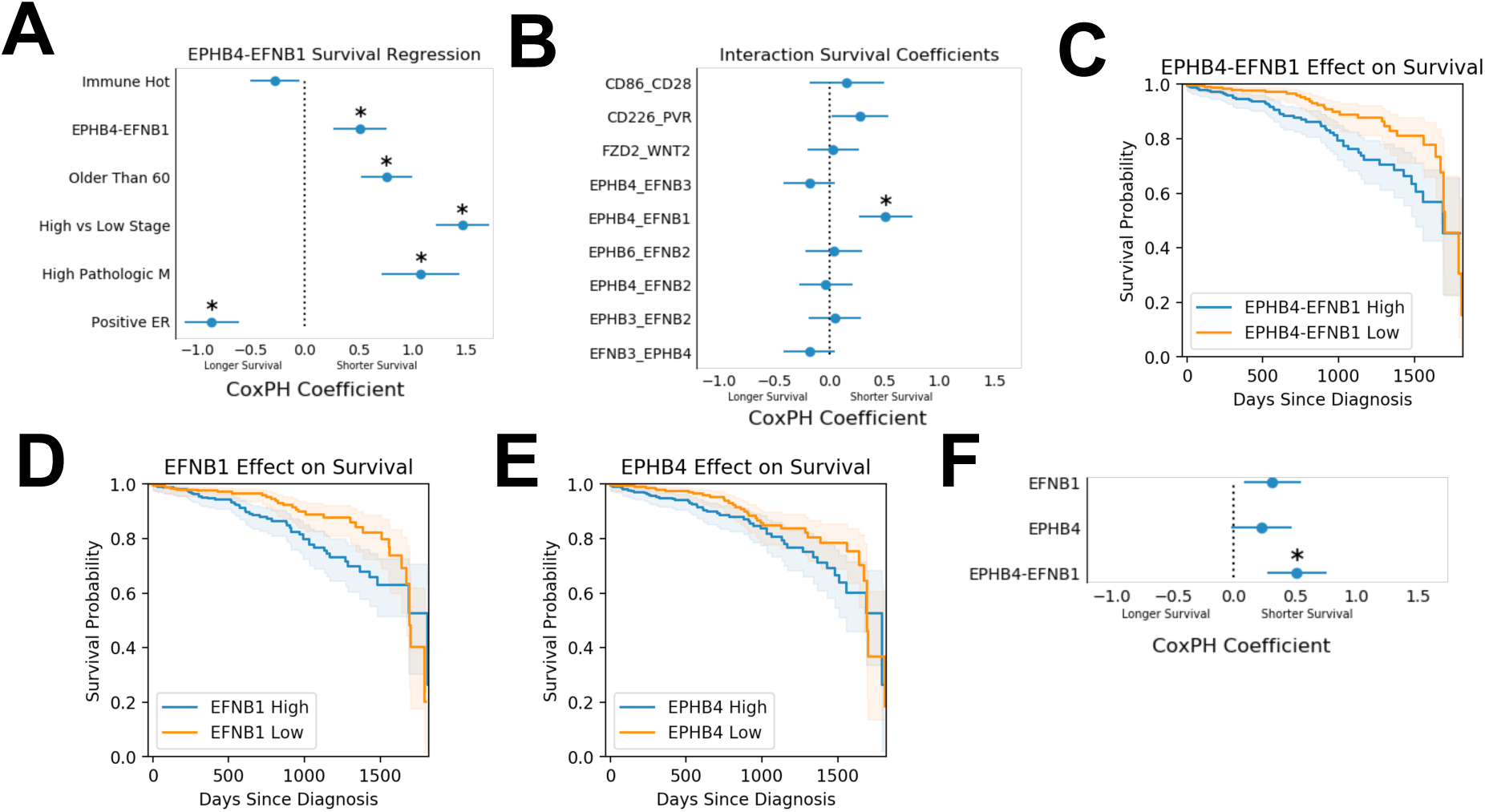
Prognostic significance of LR interactions in IDC. **(A)** Results of a multivariate Cox proportional hazards regression analysis (Hazard ratio: x-axis) including EPHB4-EFNB1 co-expression in the TCGA breast cancer cohort. Clinical covariates were selected using univariate analysis. Error bars regresent the 95% confidence interval (*p < 0.05). **(B)** Overall survival hazard ratio (x-axis) attributed to the LR pairs significant in DCIS-IDC comparison and included in 9 independent CoxPH IDC survival models. (*p=0.04). **(C-E)** Kaplan-Meier plots showing the difference in survival probability of IDC patients with high and low EPHB4-EFNB1 co-expression (C), EFNB1 (D) or EPHB4 (E) single gene expression (CoxPH p=0.04, p=0.18 and p=0.36, respectively, CoxPH). **(F)** CoxPH coefficients for separate and combined interaction models for the EPHB4-EFNB1 interaction.

## Discussion

As gene expression data is widely available for a range of diseases of the breast, we can answer more nuanced questions using comparative microenvironment modeling approaches. In particular, questions related to the molecular and functional landscape surrounding early cancer lesions and their role in cancer progression are becoming more pressing.^22,23^ We performed a meta-analysis of published gene expression studies that specifically separated the stromal and epithelial compartments in DCIS and IDC contexts. With a sufficient number of samples, we investigated functional differences using state of the art methods accounting for differences in intrinsic subtypes and cellular composition and gaining insights on the role of stromal-epithelial interactions.

Prior work has suggested modest gene expression differences between DCIS and IDC^5^, and while our observations help revisit that postulate, hierarchical clustering confirmed that in comparison to normal breast, DCIS and IDC are highly similar. The first analytical improvement we brought was to correct for cellular composition using computational deconvolution. Indeed, while the histological regions studied were microdissected, it does not mean they represent a pure population of cells. The cell-type relative abundance across samples and regions followed expected trends in cellular composition. For example in stromal regions, it is well established that TNBC and Her2 positive subtypes have higher levels of immune cells.^4,24^ In epithelial regions, infiltrating lymphocytes and macrophages, endothelial cells from increased vascularization, or fibroblasts at the invasive front can all be present.^24,25^ Similarly, DCIS lesions tended to have higher prevalence of myoepithelial cells.^26^ The computational deconvolution has limitations, especially in comparison to emerging single-cell or spatial profiling approaches. In particular, at the resolution considered, we are not able to distinguish between B- and T-lymphocytes, macrophage polarity, or fibroblast subtypes, all possibly contributing to invasion and immune escape. On the other hand, such high resolution assays have not yet been conducted on large cohorts and, for single-cell RNA-seq, are critically dependent on the availability of fresh specimens, which are impossible to get for most pure DCIS cases.

Akin to gene expression comparison, gene set enrichment analysis have been also conducted between DCIS and IDC gene expression datasets, including by the very studies that were sourced to assemble our meta-analysis. Correcting for cellular abundance, even at limited resolution, helped identify intrinsic differences in pathway activities, in contrast to differences due to variable cellular composition which may have driven the expression differences observed in previous studies. Notably Lee et al^6^ and Knudsen et al^27^ previously identified progression-associated expression patterns of genes expressed in the vasculature. Our results do not identify these same trends, likely due to our cell abundance normalization procedure. Knudsen et al additionally identified genes from a myoepithelial signature as significantly higher in invasive disease. Instead, we observed lower levels of myoepithelial gene expression in IDC as a result of cell abundance correction. In agreement with prior studies, processes linked to proliferation and ECM were dysregulated in IDC. Indeed, these processes are likely dominant in the epithelial compartment, which is the least affected by the cell abundance correction we implemented.

Thanks to the aggregation of multiple datasets and the resulting large number of samples, we were able to distinguish dysegulated processes by subtypes, revealing compelling contrasts in both epithelial and stromal functional states. The most prominent finding in the epithelium was that the majority of subtype-dependent dysregulation of cell cycle and DNA repair occurred in the ER- and Her2- context, and by extension is likely driven by the triple negative subtype. Estrogen receptor status has been associated with DNA repair capacity,^28^ but it is interesting to observe that this dependency may be limited to IDC and that proliferation and DNA repair may become increasingly dysregulated as the disease progresses. As a corollary, it may suggest that ER- and/or Her2- DCIS may not be as proliferative and genetically unstable as their invasive counterparts. Interestingly, telomere maintenance was also specifically induced in IDC in the ER- context, consistent with the evidence that altered telomere maintenance associated with invasion and stemness.^29,30^ Our results suggest that the dysregulation of this process is more prominent in IDC than DCIS.

In the stroma, we identify increased Notch or Rho GTP-ase signaling in DCIS, irrespective of subtypes. Notch signaling has been implicated in both breast cancer progression and inhibition^31^ as well as in tumor-stroma crosstalk.^32^ Notch signaling in stromal cells (fibroblasts or immune cells) contributes to carcinogenesis and drug resistance.^33^ Such pleiotropy of Notch signaling as well as the lack of information on which Notch ligands and receptors are at play renders the interpretation of our results difficult, but it may suggest that down regulation of certain stromal Notch signals are associated with invasive disease, perhaps supporting immune escape. Subtype-specific dysregulated processes in the stroma were dominated by immune signaling, stronger in IDC in ER- context, but as mentioned before may be due to residual, or uncorrected differences in cell abundance, which would predominantly affect ER- disease, where immune-infiltration is higher.

The evaluation of ligand-receptor interactions between stroma and epithelium is the most novel aspect of our report as it required the availability of expression datasets from microdissected regions in a large enough number of samples to support a robust statistical analysis. Such discovery efforts were further aided by the restriction to well-curated pairs of ligands and receptors. While much more comprehensive databases of ligand and receptor exist,^10^ they are not well validated (e.g. computationally derived) or overwhelmed with LR pairs irrelevant to breast cancer biology (e.g. neuronal communications). In contrast, we chose to restrict the analysis to experimentally proven LR pairs, for which one member was expressed in the studied dataset. While this approach is likely less sensitive, evaluating fewer pairs and relying on known biology, it facilitated the implementation of an empirical framework to test for the significance of the interactions via permutation and increased our confidence in the differential interaction observed. Notably, previous approaches to measure cell-cell interactions did not necessarily propose solutions for differential interaction testing, hence making it hard to compare interaction scores between samples and conditions. Importantly, conducting cell abundance deconvolution prior to cell-cell interaction scoring can help normalize heterogeneous samples and hence measure genuine changes in cell-cell interaction instead of those driven by a shift in cell composition. Similar approaches have been used in bulk expression studies,^34^ but, here again, the availability of single-cell RNA-seq could alleviate the need for such computational tricks and directly measure LR interaction from pure populations of cells as conducted by multiple methods recently published.^35–37^ As mentioned above, however, single-cell gene expression data is not available for pure DCIS specimens, and approaches like ours are needed to overcome the resulting lack of data. Emerging spatial profiling methods may soon replace or complement this approach by eliminating the imprecise and poorly scalable micro-dissection step and also by measuring the physical proximity and transcriptional state of cells in the stroma and epithelium. Spatial profiling could provide an ideal method to validate and extend our initial observation of the remodeling of cell-cell interaction.

The WNT2-FZD2 interaction was the most notably changing stromal-epithelial interaction, stronger in IDC in both SE and ES directions, suggesting reciprocal signaling of these molecules. Increased expression of Wnt family members has been shown to induce breast cancer in mouse models^38^ but this gene family is also involved in normal breast development and lactation. Prior gene expression studies have identified dysregulation of the Wnt pathway in breast cancer compared to healthy tissue^39^ and an upregulation of Wnt signaling in early carcinogenesis. Furthermore, the level of WNT2-FZD2 interaction is not different between DCIS and adjacent normal epithelium, suggesting the change occurs later in breast cancer progression or specifically affects patients diagnosed with IDC. Of the nine Wnt-Fzd family interactions investigated, this is the only pair that is significantly higher in IDC indicating that this specific exocrine Wnt signal may be relevant to the IDC phenotype.

Two immune-mediated interactions that were significantly stronger in IDC compared to DCIS-- CD28-CD86 (ES) and CD226-PVR (ES) - highlight immunological differences between DCIS and IDC. Both of these interactions are stimulatory interactions involving T-cells and/or NK cells and increase the susceptibility of cancer cells to death, hence consistent with their possible role in more advanced invasive disease. The CD28-CD86 (ES) directions and cell-type specificity suggest that T-cells in the epithelium are being stimulated by antigen presenting cells in the stroma, hence reflective of an ongoing active immune response^40^. Similarly, CD226-PVR (ES) stronger interaction in IDC is also mediated by immune cells from the epithelial compartment. Acknowledging that immune-cell abundance differences cannot be completely corrected computationally, these findings likely reflect the overall higher immunoreactive microenvironment environment observed in IDC. In general the IDC environment is more immunosuppressive and with T cells that are less primed for activation^24^. Consistently, the stronger CD86-CD28 interaction observed in IDC could mediate FOXP3+ regulatory T-cell homeostasis^41^ contributing to this immunosuppressive environment. Of note, the somewhat redundant CD80-CD28 interaction, which affects the same T cell homeostatic pathway, is not identified as significantly differential across normal, DCIS or IDC.

The most consistent result to come out of the microenvironment modeling approach is the 6 DCIS-enriched ephrin interactions. These are of particular interest to breast cancer as different ephrin signaling members have been implicated both as promoters or suppressors of tumor invasion.^17,42^ Ephrins themselves can be ligands or receptors based on their function and have several pleiotropic binding partners. These ephrin interactions represent 4 of the 49 unique ephrin interactions in CellPhoneDB. By using the normal samples as an outgroup we identified that 3 out of 4 the significantly changing ephrin interactions have significant monotonically decreasing trends from normal to DCIS to IDC suggesting a progressive erosion of ephrin interactions as the disease progresses. Remarkably, EPHB4-EFNB1 interaction was associated with poor survival in TCGA, indicating that this interaction may have prognostic value at multiple stages of the disease. The true prognostic value after a DCIS diagnosis would however need to be properly validated, ideally through independent stromal and epithelial gene or protein expression measurements. Given the slow and rare progression, such cohorts would have to be established prospectively across multiple institutions. Finally, given some of the nuanced phenotypes mediated by ephrins, it may be difficult to get precise mechanistic insights or determine of this erosion is truly causal to the disease progression in patients - or rather the results from high growth and proliferation fueled by other drivers and further experiments and validation will be required.^16,17,43^

Despite its innovative aspects, it is important to highlight some of the most important limitations of an analysis like the one presented here. First and foremost is the reliance on gene expression data. Most CCI interaction analysis rely on the expression of ligand and receptor genes, but the CCI are mediated at the protein level and the two do not always correlate. While changes in gene expression may be sufficient to capture long-term changes that affect cell identity and cell states, more transient changes may be mediated only at the protein level, or by changes in subcellular distribution or post-translational modifications, including glycosylation, which are ignored by our approach. Cell biology experiments will be required to faithfully capture the CCI changes. Furthermore, the study included cross-sectional samples from DCIS or IDC patients, and therefore may not truly capture changes that may occur longitudinally in individuals. This limitation is systematic in the vast majority of studies of DCIS, as longitudinal sampling cannot be performed. But since not all DCIS patients progress to IDC, there is increasing evidence that host specific factors contribute to DCIS progression or containment ^44,45^ and cross sectional studies cannot capture their contributions.

Despite the above limitations, the stromal-epithelial modeling approach considerably enriched the context of previous findings, characterizing better the source of expression differences between DCIS and IDC from a functional and micro-environmental viewpoint. Additional studies using spatial profiling and protein based cell-cell interaction observation will be needed to confirm the findings and determine their prognostic value or therapeutic utility to prevent breast cancer progression.

## Methods

### Preparation of the gene expression datasets

Gene expression datasets were obtained from each of the 6 studies available on NCBI Gene Expression Omnibus (GEO) database.^5,6,9,27,46,47^ The studies were selected with the following criteria: (1) gene expression profiling by Affymetrix microarray (2) laser-capture microdissection or manual macro-dissection of breast tissue specimens into stromal and epithelial compartments and (3) included samples of invasive ductal carcinoma (IDC) or ductal carcinoma in situ (DCIS). Where available, CEL files were downloaded directly from GEO, background corrected and normalized using the Robust Multi-Array Average (RMA)^48^ and multiple probes for the same gene were collapsed to the mean value. Where CEL files were unavailable the processed RMA values on GEO were used. The RMA expression levels from each of the separate datasets were combined and batch effects were corrected using ComBat from the sva package^49^ accounting for differences in tissue compartment and disease state in the design matrix (“mod” argument). The resulting batch-corrected RMA matrix was used as-is for all subsequent analyses except for the principal component analysis (PCA) and hierarchical clustering where z-scoring was performed prior to analysis. Cosine distance with complete linkage was used as the distance metric for hierarchical clustering.

### Molecular subtype classification

The “molecular.subtyping” function in the genefu package (version 2.22.1) was used to determine the PAM50 intrinsic subtypes for the epithelial regions of DCIS and IDC biopsies from normalized expression values. Her2 status and ER status were assigned based on the expression of the ERBB2 (z>1 is positive) and ESR1 genes (z>0 is positive), respectively.

### Accounting for cellular composition

#### Cell abundance measurement

CIBERSORTx^13^ was used to estimate cell abundance from the normalized expression values. The reference dataset was derived from single-cell RNA-seq data from 28 normal breast reduction mammoplasty donors for a total of 86,136 cells assigned to 10 cell type clusters. Histologically related clusters were aggregated into 6 major cell lineages: fibroblasts, myoepithelial cells, luminal cells, lymphocytes, macrophages and vascular and endothelial cells. The CIBERSORTx HiRes Docker (release date Sep 2019)^50^ was used in S-mode to quantify cell abundance from the batch-corrected expression matrix, *G*_*IxJ*_, with genes *I* and samples *J*, using the cell lineages *C*, identified in the single-cell RNA-seq data. This method estimates a matrix of cell lineage specific expression, *A*^*j*^_*IxC*_, and a vector of cell lineage abundance, *B*^*j*^_*C*_, according to the following equation for every sample *j* in *J*: *G*_*Ixj*_ *= A*^*j*^_*IxC*_ *B*^*j*^_*C*_. CIBERSORTx uses the marker genes for each cell lineage in the single-cell RNA-seq data to constrain this equation and identify bulk gene expression variation that can be attributed to each of the cell lineages provided..

#### Normalization of the gene expression matrix

The gene expression matrix was normalized to account for differences in cell abundance between samples. First, a median cell abundance vector for each tissue compartment is computed by taking the median of all cell abundance vectors within stromal and epithelial samples separately giving *B*^*str*^_*C*_ and *B*^*epi*^_*C*_. We expect the stromal and epithelial compartments to have different cell composition so we normalize them separately. Then the *G*_*IxJ*_ matrix is normalized for cell abundance differences within each tissue compartment giving *G*^*N*^_*IxJ*_ according to the following procedure: if a sample, *j*, is from the stromal compartment then *G*^*N*^_*Ixj*_ *= A*^*j*^_*IxC*_ *B*^*str*^_*C*_, otherwise: *G*^*N*^_*Ixj*_ *= A*^*j*^_*IxC*_ *B*^*epi*^_*C*_ where *A*^*j*^_*I,C*_ is the corresponding cell lineage specific expression matrix for sample *j*. Finally, *G*^*N*^_*IxJ*_ is log-transformed to return to RMA space.

### Gene Set Enrichment Analysis

MSigDB Hallmarks (H) and REACTOME (C2 REACTOME) collections were used in all gene set centric analyses. Gene set enrichment analysis (GSEA) was performed using cell-abundance normalized expression values from G^N^_i,j_. The GSEA implementation in *gseapy (https://github.com/zqfang/GSEApy)* was used to identify gene sets significantly changing between DCIS and IDC for each subtype and histological region considered. Gene sets with a minimum of 15 genes that were detected in the dataset were retained for permutation analysis. The significance was determined using a permutation test on the phenotype labels for a total of 10,000 permutations. Gene sets with an FDR of less than 0.01 were considered significant.

### Differential Cell-Cell Interaction Measurement

The CellPhoneDB ligand-receptor (LR) database (N=279 pairs) was filtered for detected interactions between ligands and receptors in the cohort, resulting in 139 ligand-receptor pairs. Two separate ligand-receptor interaction models were considered: one for stromal-epithelial (SE) and another for epithelial-stromal (ES). For each LR pair in the ES model, the interaction score was computed from the outer sum of the ligand RMA expression in the epithelium and the receptor RMA expression in the stroma (and vice-versa for the SE model). The median score among DCIS-DCIS sample pairs and IDC-IDC sample pairs were compared. The significance of the observed DCIS-IDC difference was determined in comparison to a null distribution of interaction scores differences obtained from 10,000 permutations randomly shuffling within epithelial and stroma regions. Two-sided p-values were computed from this null distribution and corrected for multiple hypothesis testing using the Benjamini-Hochberg procedure. Interactions with an empirical FDR of less than 0.1 were considered significant. The same analytical workflow was used to compare IDC to normal and DCIS to normal.

### Survival Analysis in TCGA Breast Cancer Cohort

The PanCan TCGA gene expression data RNA-seq V2.0 was downloaded from the PanCan GDC portal. Clinical annotations from the same website were used. Univariate Cox proportional hazards (CoxPH) models were used to select important clinical features for the final multivariate model. Analyses were performed in Python 3.9.12 using the lifelines package version 0.27.0. Hormone receptor results were only used if the sample was labeled “Positive” or “Negative”. T, N and M pathological staging values were simplified to remove sub-stages (e.g. T1a becomes T1), and age was thresholded at 60 years old in line with previous studies.^51^ The multivariate CoxPH model included only clinical variables that were significant in univariate analyses. For each ligand-receptor pair tested the interaction score was derived from the product of RSEM values and binarized into “Low” and “High” groups according to the median interaction score.

## List of abbreviations

DCIS: Ductal Carcinoma In Situ
IDC: Invasive Ductal Carcinoma
LR: Ligand-Receptor
ECM: Extracellular Matrix
CCI: Cell-Cell Interaction
SE: Stromal-Epithelial
ES: Epithelial-Stromal
RSEM: RNA-seq by Expectation-Maximization
TCGA: The Cancer Genome Atlas
FDR: False Discovery Rate

## Declarations

### Ethics approval and consent to participate

The study was performed on existing public data and the need for consent was waived by the UCSD IRB.

### Consent for Publication

Not applicable

## Acknowledgements

We thank Erick Armingol Gonzalez and Nathan Lewis for helpful discussions.

## Availability of Data and Materials

All gene expression dataset were obtained from the NCBI Gene Expression Omnibus. The microdissected IDC, DCIS and normal datasets are available under the following accession numbers: GSE41194, GSE41196, GSE41197, GSE41227, GSE41198, GSE14548, GSE16873, GSE24506, GSE33692, GSE3893. The single-cell RNA-seq data is available under GSE198732.

## Authors’ contributions

A.O. A.D. performed the analysis, L.M., Z.G., C.Y., P.T., contributed analysis and interpretation of the data. O.H. directed the study. A.O., O.H, wrote the manuscript. All authors reviewed and approved the manuscript.

